# Heterogeneities in infection outcomes across species: sex and tissue differences in virus susceptibility

**DOI:** 10.1101/2022.11.01.514663

**Authors:** Katherine E Roberts, Ben Longdon

## Abstract

Species vary in their susceptibility to pathogens, and this can alter the ability of a pathogen to infect a novel host. However, many factors can generate heterogeneity in infection outcomes, obscuring our ability to understand pathogen emergence. Such heterogeneities can alter the consistency of responses across individuals and host species. For example, sexual dimorphism in susceptibility means males are often intrinsically more susceptible than females (although this can vary by host and pathogen). Further, we know little about whether the tissues infected by a pathogen in one host are the same in another species, and how this relates to the harm a pathogen does to its host. Here, we first take a comparative approach to examine sex differences in susceptibility across 31 species of Drosophilidae infected with Drosophila C Virus (DCV). We found a strong positive inter-specific correlation in viral load between males and females, with a close to 1:1 relationship, suggesting that susceptibility to DCV across species is not sex specific. Next, we made comparisons of the tissue tropism of DCV across seven species of fly. We found differences in viral load between the tissues of the seven host species, but no evidence of tissues showing different patterns of susceptibility in different host species. We conclude that, in this system, patterns of viral infectivity across host species are robust between males and females, and susceptibility in a given host is general across tissue types.

## Introduction

Emerging pathogens often arise from a host shift event – where a pathogen jumps into and establishes in a novel host species. Species vary in their susceptibility to pathogens, but little is known about the factors underlying this variation, and whether differences between clades are due to the same or different factors [1, 2]. Understanding this is critical for determining which hosts pathogens are likely to jump between, and the harm they cause to their hosts. The host phylogeny has been shown to be an important determinant of host shifts in a range of systems [3–8] as well as being important for understanding how pathogen virulence may change when a pathogen finds itself in a new host [9, 10]. For example, virulence tends to increase, and onward transmission and pathogen load decrease, with greater evolutionary distance between donor and recipient hosts [10–12]. In addition, clades of closely related species tend to have similar levels of susceptibility independent from their distance to the pathogens natural host [4, 9].

However, such patterns of susceptibility across species can be affected by heterogeneities in infection outcomes. These can be viewed at the level of the individual heterogeneities within species, and by looking at whether these heterogeneities are consistent or different across host species. Experimental studies typically try and minimise within species or environmental effects. Often overlooked is variation arising from sexual dimorphism, as typically only one sex is utilized to remove between sex differences [13]. Sexual dimorphism is seen across most animal systems in a range of life history traits from body size, growth rate, reproductive effort and immunity [14–16]. In mammals, males and females often differ in their pathogen burdens and mortality rates [17]. For example, in SARS-COV-2 infection in humans, women have a lower risk of morbidity and mortality than men [18]. In HIV infected individuals, women have up to 40% lower HIV viral RNA in circulation but a greater likelihood of developing AIDS than men with matched viral loads [19]. Sexually transmitted infections – which are primarily transmitted between the sexes – are particularly prone to sex biased infection through either exposure differences or sex biased virulence [20–22]. Sex biases in parasitism rates in mammals have been suggested to be due to males investing in traits that favour their reproductive success, which trade off against somatic maintenance, including immunity. In support of this, sex biased parasitism is positively correlated with sexual size dimorphism [17]. In insects a comparative analysis found the degree of sex biased parasitism and mortality is explained by an interaction between the mating system (polygynous vs non-polygynous) and sexual size dimorphism [23]. This difference is consistent with parasites having a greater impact on the survival of male insects compared to females (particularly in polygynous species where males are larger than females). Other hypotheses suggest that as longevity is a major determinant of female fitness, investment in costly immune responses are more important than for males, who can maximise their fitness with shorter term mating success [24]. As such, investment in reproduction may trade-off with immunity in different ways between males and females. There is some support for this hypothesis in mammals but a lack of supporting data for insects [15]. However, many experimental studies of host-parasite interactions do not compare differences between sexes [13]. Furthermore, for most pathogens we have little understanding of whether sex differences are consistent across host species, which has important implications for our understanding of pathogen emergence.

Despite phylogenetic patterns of host susceptibility having been observed in a range of systems [3, 6–8], we know little about why species vary in their susceptibilities. For example, given equal exposure why do we see high mortality in some species but little in others? One factor that appears important in determining the severity of disease is the tissue tropism of a pathogen. In humans, RNA viruses with neural tropism or generalised systemic tropism tend to result in severe disease [25]. In terms of the patterns of susceptibility across species, virulence may be a consequence of a virus infecting a sensitive tissue or organ resulting in damage by the pathogen directly or by autoimmunity. For example, in bacterial meningitis the pathology is a consequence of bacteria infecting the cerebrospinal fluid and resulting in inflammatory autoimmune damage to the central nervous system [26]. Alternatively, it may be due to the pathogen getting into a particularly permissive tissue type and proliferating to high levels. Virus macroevolutionary change is thought to be driven by cross-species transmission or codivergence rather than by acquiring new niches – or tissues – within a host [27]. Likewise, the host specificities of viruses are thought to be more labile than tissue specificities [28]. As such, heterogeneity in the tissues infected may explain variation in susceptibility across host species.

Here we use a *Drosophila-virus* system to examine the factors underlying susceptibility across host species. Across *Drosophila* species many physiological traits show sexual differentiation [14]. Using a comparative approach, we firstly ask if the patterns of infection seen across the host phylogeny [3, 5, 9, 29] differ between males and females. We infected both males and females of a panel of 31 species of Drosophilidae with Drosophila C Virus (DCV), a positive sense RNA virus in the family Dicistroviridae. Males of *Drosophila melanogaster* have previously been reported to have higher viral loads than females [30] and show greater rates of shedding, lower clearance and higher transmission potential of DCV, although these traits can interact with host genotype [31]. Viral load has shown to have a strong positive correlation with mortality across host species [9, 29]. DCV has been reported to show tissue tropism in *D. melanogaster*, with high levels of infection in the heart tissue, fat body, visceral muscle cells around the gut (midgut) and food storage organ (crop) [32, 33]. To test if the same patterns of tissue infection were observed across species, we then made comparisons of the tissue tropism of DCV in 7 species of fly.

## Methods

### Viral Infections

Thirty one species of Drosophilidae were used to examine sex differences in viral infection. Stock populations were reared in the laboratory in multi generation populations, in Drosophila stock bottles (Fisherbrand) on 50 ml of their respective food medium (Table S1) at 22°C and 70% relative humidity with a 12-hour light-dark cycle. Flies were then collected twice a day in order to try and control for age of maturity in an effort to minimize the chances that flies would have reached sexual maturity and mated before the sexes were separated out. Although no effect of mating status on DCV viral load was previously observed in *D. melanogaster*, this can vary by host genotype and mating status is known to affect susceptibility to other pathogens [31].

To examine differences in viral load between males and females, two vials of 0-1 day old males flies and two vials of 0-1 day old female flies were collected daily for each species. Flies were tipped onto fresh vials of food every day to minimise differences in the microbiomes of flies (Broderick & Lemaitre, 2012; Blum *et al.*, 2013). All vials were kept for 10 days in order to check for larvae, as a sign of successful mating. Only 4 vials from 3 species were found to contain larvae, these were one vial of *D. sturtevanti*, two vials of *Scaptodrosophila lativittata* and one vial of *Zaprionus tuberculatus*. After 3 days flies were experimentally infected with DCV. Three replicate blocks were carried out, with each block being completed over consecutive days. The order of experimental infection was randomized each day so that both sex and species were randomised. We carried out three biological replicates for each species for each sex at time zero and 2 days post infection. There was a mean of 17 flies per replicate (range across species = 12-20).

Viral challenge was carried out by needle inoculation of Drosophila C virus (DCV) strain B6A [34], derived from an isolate collected from *D. melanogaster* in Charolles, France [35]. The virus was prepared as described previously [36]. DCV was grown in Schneider’s Drosophila line 2 cells and the Tissue Culture Infective Dose 50 (TCID50) per ml was calculated using the Reed-Muench end-point method. Flies were anesthetized on CO2 and inoculated using a 0.0125 mm diameter stainless steel needle bent to a right angle ~0.25mm from the end (Fine Science Tools, CA, USA). The bent tip of the needle was dipped into the DCV solution (TCID50 = 6.32×10^9^) and pricked into the anepisternal cleft in the thorax of the flies [9, 37]. This mode of infection is used as it creates a more reproducible infection that oral inoculation, which is found to cause stochastic infection outcomes in *D. melanogaster* [32]. Both methods of infection have been shown to produce systemic infections with the same tissues ultimately becoming infected [32].

To control for relative viral dose between species a time point zero sample of one vial of flies was immediately snap frozen in liquid nitrogen as soon as they were inoculated. The second vial of flies were inoculated and placed onto a fresh vial of food, and returned to the incubator. Two days after challenge (+/- 1 hour) these flies were snap frozen in liquid nitrogen. This time point is chosen as the sampling time point as previous studies show a clear increase in viral growth but little mortality at this point in infection [5, 29]. Each experimental block contained a day 0 and day 2 replicate for each sex and each species (31 species × 2 sexes × 3 experimental blocks).

### Measuring the change in viral load

Using quantitative Reverse Transcription PCR (qRT-PCR) we measured the change in viral load in male and female flies from day 0 to day 2 post-infection. Total RNA was extracted from the snap frozen flies by homogenizing them in Trizol reagent (Invitrogen) using a bead homogenizer for 2 pulses of 10 seconds (Bead Ruptor 24; Omni international) and stored at −70°C for later extraction. Samples were defrosted and RNA extracted as described previously [29]. Briefly, Trizol homogenized flies were processed in a chloroform isopropanol extraction, eluted in water and reverse-transcribed with Promega GoScript reverse transcriptase (Sigma) and random hexamer primers. Quantification of the change in viral RNA load was calculated in relation to a host endogenous control, the housekeeping gene *RpL32*. Primers were designed to match the homologous sequence for each of the experimental species that crossed an intron– exon boundary so will only amplify mRNA. qRT-PCR was carried out on 1:10 diluted cDNA using Sensifast Hi-Rox Sybr kit (Bioline). Two qRT-PCR reactions (technical replicates) were carried out per sample with both the viral and endogenous control primers. All melt curves were checked to verify that the correct products were being amplified. All experimental plates had experimental replicates distributed across the plates in a randomized block design to control for between plate differences. Each qRT-PCR plate contained three standard samples. A linear model which included plate ID and biological replicate ID was used to correct the cycle threshold (C_t_) values between plates. Any technical replicates that had Ct values more than two cycles apart after the plate correction were repeated. Change in viral load was calculated as the mean C_t_ value of the pairs of technical replicates. We then used these to calculate the ΔC_t_ as the difference between the cycle thresholds of the viral DCV qRT-PCR and the RpL32 endogenous control for each sample. The Ct of the day 2 flies relative to day 0 flies was then calculated as, 2^-ΔΔC_t_^; where ΔΔC_t_ = ΔC_t_ day0 – ΔC_t_ day2.

### Body Size

We measured wing size of the flies to control for between species and sex differences in body size. In Drosophilidae wing length has been shown to be a good proxy for body size (Huey *et al.*, 2006). For measurement wings were removed from a mean of 15 male and females flies of each species (range 10–18), stored in 80% ethanol, and later photographed under a dissecting microscope. The length of the IV longitudinal vein from the tip of the proximal segment to where the distal segment joins vein V was recorded Using ImageJ software (version 1.48), and the mean taken for each sex of each species.

### Inferring the host phylogeny

We used a previously inferred phylogenetic tree [5] using seven genes (mitochondrial; *COI, COII*, ribosomal; *28S* and nuclear; *Adh, SOD, Amyrel, RpL32*). Briefly, we downloaded publicly available sequences from Genbank and where these were not available they were Sanger sequenced from our laboratory stocks. For each gene the sequences were aligned in Geneious (version 9.1.8, www.geneious.com) [38] using the global alignment setting, with free end gaps and 70% similarity IUB cost matrix. The phylogeny was inferred using these genes and the BEAST programme (v1.10.4) [39]. Genes were partitioned into three groups; mitochondria, ribosomal and nuclear, each with separate relaxed uncorrelated lognormal molecular clock models using random starting trees. Each of the partitions used a HKY substitution model with a gamma distribution of rate variation with 4 categories and estimated base frequencies. Additionally, the mitochondrial and nuclear data sets were partitioned into codon positions 1+2 and 3, with unlinked substitution rates and base frequencies across codon positions. The tree-shape prior was set to a birth-death process.

We ran the BEAST analysis three times to ensure convergence for 1000 million MCMC generations sampled every 10000 steps. On completion the MCMC process was examined by evaluating the model trace files using the program Tracer (version 1.7.1) (Rambaut *et al.*, 2014) to ensure convergence and adequate sampling. The consensus constructed tree was then visualised using FigTree (v1.4.4) (Rambaut, 2006).

### Tissue Tropism

In order to examine patterns of tissue infection across species we infected 7 species of flies used above; *D. melanogaster, D. stuventi, S. lativaitata, D. pseudooscura, D. virilis, D. prosaltans* and *D. littoralis*). Male flies were infected with DCV using the same inoculation method as described above. Two days post infection flies were placed on ice to sedate them, they were then surface sterilized in ice-cold 70% ethanol before being dissected. The head, crop, gut (all parts), malpighian tubules, sex organs (testis and accessory glands) and abdominal cuticle including the attached fat body (hereafter referred to as body) were dissected from each male fly and placed into individual tubes on ice. Six individual flies were pooled per replicate and then snap frozen in liquid nitrogen for later RNA extraction. For each species there were six replicate pools of each of the six tissue types. At the same time as the dissections were carried out whole flies were snap frozen for a “whole fly” comparative viral load measure. All samples were processed as per the methods for viral load quantification as described above.

### Statistical analysis

#### Sex differences

Viral load in males and females were analysed using phylogenetic mixed models. We fitted all models using a Bayesian approach in the R package MCMCglmm [40, 41]. We used a multivariate model with viral load of each sex as the response variable.

The models took the form of:

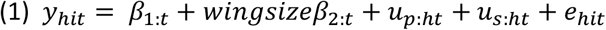

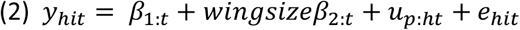

where y is the change in viral load of the *i*th biological replicate of host species *h*, for trait *t* (male or female). *β* are the fixed effects, with *β*_1_ being the intercepts for each trait and *β*_2_ being the effect of wing size. *u_p_* are the random phylogenetic species effects, and *e* the model residuals. Models were also run which included the mating status of the species as a fixed effect. We included this as a binary response for any species that had offspring in at least one replicate vial (we only found evidence of mating for three species: *D. sturtevanti, S. lativittata* and *Z. tuberculatus*). Model (1) also includes a species-specific component independent of the phylogeny *u _s:ht_* that allow us to estimate the proportion of variation that is not explained by the host phylogeny *v_s_* (Longdon et al., 2011). However, this was removed from model (2) as model (1) failed to separate the phylogenetic and species - specific effects. The main model therefore assumes a Brownian motion model of evolution (Felsenstein, 1973). The random effects and the residuals are assumed to follow a multivariate normal distribution with a zero mean and a covariance structure **V_p_** ⊗ **A** for the phylogenetic affects and **V_e_** ⊗ **I** for the residuals, **V_s_** ⊗ **I** for species-specific effects, (⊗ here represents the Kronecker product). **A** is the phylogenetic relatedness matrix, **I** is an identity matrix and the **V** are 2×2 (co)variance matrices describing the (co)variances between viral load of the two sexes. The phylogenetic covariance matrix, **V_p_,** describes the phylogenetic inter-specific variances in each trait and the inter-specific covariances between them, **V_s_,** the non-phylogenetic between-species variances. The residual covariance matrix, **V_e_,** describes the within-species variance that can be both due to real within-species effects and measurement or experimental errors. The off-diagonal elements of **V_e_** (the covariances) are not estimable because each vial only contains one sex and therefore no vial has multiple measurements, so were set to zero. The MCMC chain was run for 1,300 million iterations with a burn-in of 30 million iterations and a thinning interval of 1 million. All the models were run with different prior structures (as in [5]) in order to test results for sensitivity to the use of priors, but note they all gave similar results.

The proportion of between species variance that can be explained by the phylogeny was calculated from model (1) using the equation 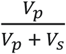, where *V_p_* and *V_s_* represent the phylogenetic and species specific components of between-species variance respectively, and is equivalent to phylogenetic heritability or Pagel’s lambda [42, 43]. The repeatability of susceptibility measurements was calculated from model (2) as 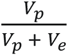, where *V_e_* is the residual variance. Inter-species correlations in viral load between each method were calculated from model (2) *V_p_* matrix as 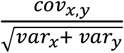 and the slopes (*β*) of each relationship as 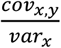. Parameter estimates stated below are means of the posterior density, and 95% credible intervals reported (CIs) were taken to be the 95% highest posterior density intervals.

#### Tissue tropism

Viral load data across species and tissues was analysed using a linear mixed effects model using the lmer function in the lme4 package in R [41, 44] with models compared using the anova function and the resulting *P* values reported. Tissue type, species and their interaction were included as fixed effects and experimental replicate as a random effect to account for the individual pool that each set of tissues came from. With only seven species there is little power to carry out models controlling for phylogeny which is why species was fitted as a fixed effect.

### Data availability

All data and scripts are available at dx.doi.org/10.6084/m9.figshare.21437223.

## Results

### Sex differences in viral load

To examine if the sexes respond the same way to viral infection we infected 31 species of Drosophilidae with DCV and quantified the change in viral load at 2 days post infection using qRT-PCR. In total we infected 6324 flies across 186 biological replicates (biological replicate = change in viral load from day 0 to day 2 post-infection), with a mean of 17 flies per replicate (range across species = 12-20).

The mean change in viral load across all species was similar between the sexes (females = 12.59, 95% CI = 1.16, 23.80; males = 12.93, 95% CI = −0.65, 26.25). We found strong positive interspecific correlation between the viral load of females and males (correlation = 0.92, 95% CI = 0.78, 1.00; Figure 1). The estimate of the slope is close to 1 (*β* = 0.99, 95% CI= 0.58, 1.38) suggesting males and females respond similarly to infection.

**Figure 1.**
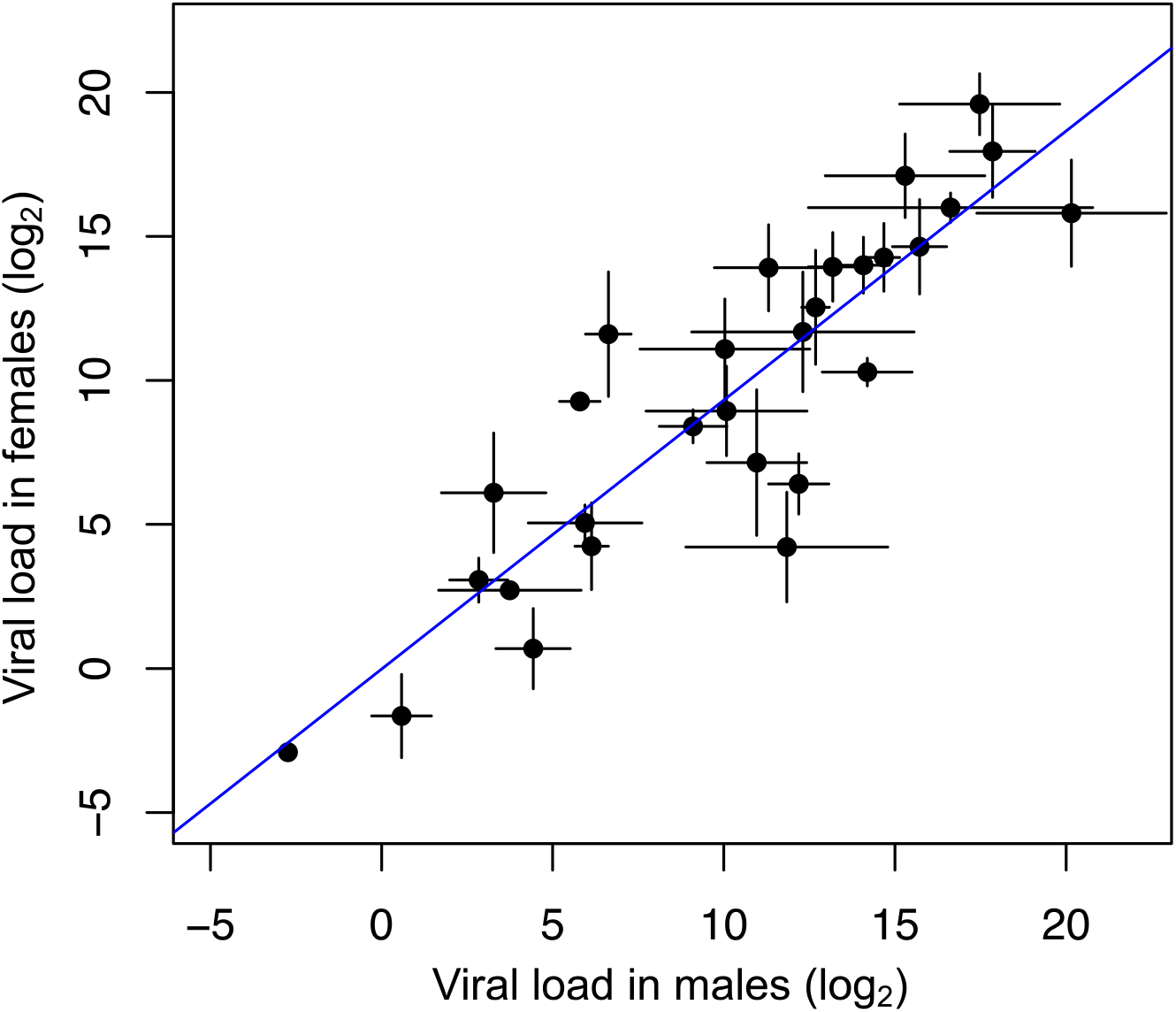
Correlation between viral load in males and females. Each point represents a species mean, error bars show standard errors and the trend line is estimated from a linear model.

The full model including the species-specific random effect independent of the host phylogeny (*u _s:ht_*) allowed us to calculate the proportion of the variation between the species that can be explained by the phylogeny 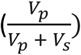, equivalent to phylogenetic heritability or Pagel’s lambda [42, 43]. The host phylogeny explains a large proportion of the inter-specific variation for both males and females (females = 0.68, 95% CIs: 0.06, 0.99; males = 0.66, 95% CIs: 0.04, 0.99) consistent with previous findings for males [3, 5, 9, 29]. However, we note these estimates have broad confidence intervals, due to the model struggling to separate out the phylogenetic and non-phylogenetic components. The repeatability of viral load across species was relatively high for both sexes (females = 0.63, 95% CIs = 0.41, 0.80; males = 0.52, 95% CIs = 0.31, 0.74). We found no effect of either body size (−0.05, 95% CI’s = −0.28, 0.19) or mating status (0.34, 95% CI’s = −5.63, 6.27) on viral load.

### Tissue Tropism

To look at the how the tissue tropism of DCV varied across host species, we infected seven species of fly with DCV and dissected them into six tissue types. We found large effects of species on viral load (χ^2^= 320.65, d.f=6, *P*<0.001) with >18 million fold difference in viral load between the least and most susceptible species. Tissues differed in their viral loads to a lesser extent, with the maximum difference being seen in *D. pseudobscura* with an approximately 550 fold difference in viral load between the least and most susceptible tissues (χ^2^= 15.264, d.f=4, *P*=0.009). There was no evidence of tissues showing different patterns of susceptibility in different hosts i.e. no evidence for a tissue-by-species interaction (χ^2^=41.515, d.f=30, *P*=0.079).

## Discussion

Here, we examined whether heterogeneities in infection outcomes altered patterns of susceptibility across host species. We first examined whether males and females responded in consistent or different ways to infection with DCV. We found that viral susceptibility between females and males of 31 host species showed a strong positive correlation with a close to 1:1 relationship, suggesting that susceptibility across species is not sex specific. We next examined whether heterogeneities in the tissues infected across host species altered the outcome of infection. We found differences in viral load between tissues of seven host species, but no evidence of tissues showing different patterns of susceptibility in different host species.

A difference between the sexes in immune function and resistance has been in found in a range of studies [15, 45]. However, this is not universal, and interactions between the host, pathogen and environmental factors can alter the outcome of infections [46]. Previous meta-analyses have found mixed results. For example, some studies of arthropods have found little evidence for consistent sex differences in parasite prevalence or intensity [47] (although these were largely from natural infections which may have inherently greater sources of variation). Likewise, a phylogenetically controlled meta-analysis of 105 species (including 30 insect species) of immune responses found no evidence for sex biases [48]. Other studies have reported a small male bias in parasitism rates for polygynous insects, but a significant female bias for non-polygynous species, with the extent of sex bias parasitism increasing with the degree of sexual size dimorphism [23]. However, other studies of insects (with a smaller number of species) have reported sex differences in some immune traits [15, 23] and sex biased prevalence and impacts are well known for sexually transmitted infections in insects [20, 21]. In mammals, males have been shown to often have greater pathogen burdens, with parasitism rates positively correlated to male biased sexual size dimorphism [15–17, 23].

In *Drosophila melanogaster*, some previous studies have reported males have higher DCV viral loads than females [30]. However, others found no effect of sex on viral load, but did find effects of sex on viral shedding, clearance, and transmission potential, with these traits showing interactions with host genotype [31]. Sexual dimorphism in infection avoidance behaviour has also been reported, when female flies previously exposed to DCV were found to prefer a clean food source indicating a potentially important dimorphism in infection avoidance [49]. Here, we used controlled experimental conditions, but in nature sex differences in behaviour, or how the sexes interact with the environment may lead to differences in pathogen load. A caveat is that the flies used here were of a fixed age and largely virgins – future studies should explore if age and mating status can affect these results.

The tissue tropism results here show that susceptibility in a given host is general across tissue types – for example *D. sturtevanti* has a high viral load across all tissues whereas *D. virilis* has relatively low viral loads in all tissues (Figure 2). Mortality to DCV infection has previously been shown to show a strong positive correlation with viral load [9, 29]. The data presented here show the susceptibility of a given species is general across all tissue types. This does not exclude the possibility that pathology is due to high viral loads in a given tissue, but does suggest that the mechanism restricting viral load is general across tissues. This may be linked to the ability of the virus to bind to or enter hosts cells, utilise the hosts cellular components for replication or to avoid or suppress the host immune response [50]. Further work should explore this further in a range of conditions including flies of both sexes and of varying ages. Comparative studies of human viruses have identified the tissue tropism of viruses to be a significant determinant of virulence; viruses that cause systemic infections (across multiple organs) or that have neural or renal tropisms are most likely to cause severe virulence [25]. It has been suggested that high levels of non-adaptive virulence can be the result of pathogens infecting tissues that do not contribute to onward transmission [26]. Other studies have shown differences in host physiology can be important in determining the virulence of a novel pathogen [10]. However, further understanding of how infection results in pathology (i.e. in which tissue the disease tropism occurs [51]) and how virulence is correlated with transmission potential in infections, is needed to explore this further.

**Figure 2:**
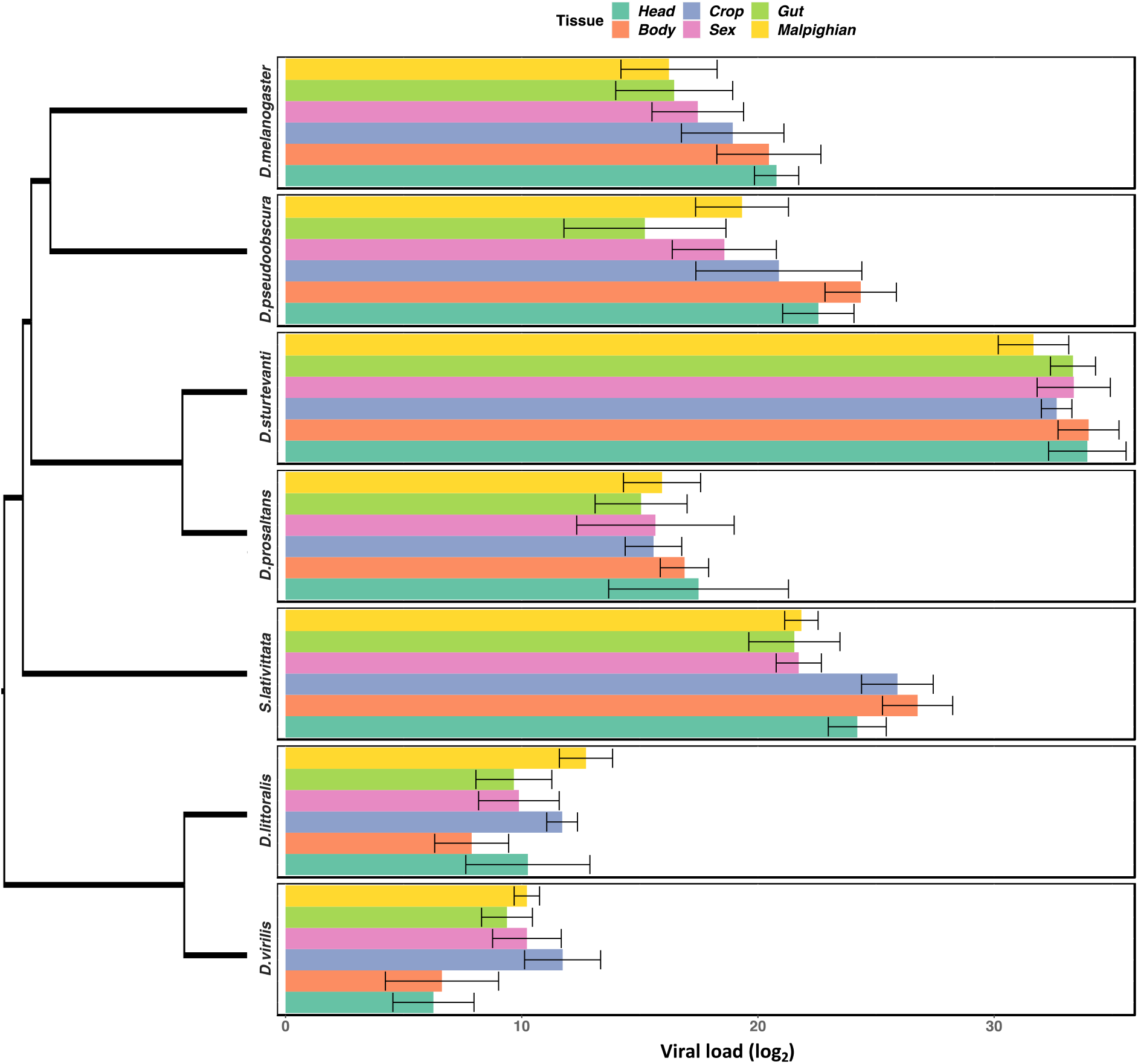
Tissue tropism data from DCV infected flies. **Male** flies were dissected into the six tissue types 2 days post-infection before undergoing RNA extraction and quantification of viral load. On the right is the phylogeny of the host species. The bars of the individual panels are organised following the order of the *D. melanogaster* ranked from the tissue with lowest to highest viral load. Error bars show standard errors.

In summary, our results show little evidence for sexual dimorphism in susceptibility to viral infection across species. As such susceptibility in one sex is predictive of that in the other. We find that susceptibility of a species is general across tissue types, suggesting virulence is not due to species specific differences in viral tropism. The patterns of susceptibility observed across species do not appear to be affected by heterogeneities in sex or tissue tropism. Further work is needed to explore how sex differences can vary with factors such as host age, mating status, the environment and pathogen type, and the underlying mechanisms as to why species vary in their susceptibility.

## Acknowledgments

Thanks to group members for discussions of these data. For the purpose of Open Access, the author has applied a CCBY public copyright licence to any Author Accepted Manuscript version arising from this submission. Many thanks for Alison Duncan, Greg Hurst and an anonymous reviewer for useful comments that improved the manuscript. This preprint (v2) has been reviewed and recommended by Peer Community in Evolutionary Biology https://doi.org/10.24072/pci.evolbiol.100638

## Supplementary information

**Table S1:** Full list of species used in the sex difference experiment and their rearing food for stock populations. All cornmeal and proprionic medium have dried yeast sprinkled onto the surface of the food, other food types do not unless stated below. The recipes for the food types are described here https://doi.org/10.6084/m9.figshare.21590724.v1

| Species | Food |
| --- | --- |
| *D.affinis* | Malt |
| *D.americana* | Malt |
| *D.ananassae* | Cornmeal |
| *D.arizonae* | Banana |
| *D.buzzatii* | Malt |
| *D.erecta* | Malt + yeast |
| *D.flavomontana* | Malt + yeast |
| *D.hydei* | Cornmeal |
| *D.immigrans* | Malt + yeast |
| *D.lacicola* | Malt |
| *D.littoralis* | Banana |
| *D.mauritiana* | Proprionic |
| *D.melanogaster* | Cornmeal |
| *D.montana* | Malt + yeast |
| *D.novamexicana* | Banana |
| *D.obscura* | Proprionic |
| *D.persimilis* | Malt |
| *D.prosaltans* | Proprionic |
| *D.pseudoobscura* | Malt |
| *D.putrida* | Propionic |
| *D.santomea* | Cornmeal |
| *D.sturtevantii* | Cornmeal |
| *D.takahashii* | Cornmeal |
| *D.teissieri* | Cornmeal |
| *D.virilis* | Propionic |
| *H.duncani* | Propionic |
| *S. lativittata* | Banana |
| *S.lebanonensis* | Propionic |
| *Z. inermis* | Banana |
| *Z. taronus* | Banana |
| *Z. tuberculatus* | Banana |

